# Enhanced competitive protein exchange at the nano-bio interface enables ultra-deep coverage of the human plasma proteome

**DOI:** 10.1101/2022.01.08.475439

**Authors:** Daniel Hornburg, Shadi Ferdosi, Moaraj Hasan, Behzad Tangeysh, Tristan R. Brown, Tianyu Wang, Eltaher M. Elgierari, Xiaoyan Zhao, Amir Alavi, Jessica Chu, Mike Figa, Wei Tao, Jian Wang, Martin Goldberg, Hongwei Xia, Craig Stolarczyk, Serafim Batzoglou, Asim Siddiqui, Omid C. Farokhzad

## Abstract

We have developed a scalable system that leverages protein-nano interactions to overcome current limitations of deep plasma proteomics in large cohorts. Introducing proprietary engineered nanoparticles (NPs) into a biofluid such as blood plasma leads to the formation of a selective and reproducible protein corona at the particle-protein interface, driven by the relationship between protein-NP affinity and protein abundance. Here we demonstrate the importance of tuning the protein to NP-surface ratio (P/NP), which determines the competition between proteins for binding. We demonstrate how optimized P/NP ratio affects protein corona composition, ultimately enhancing performance of a fully automated NP-based deep proteomic workflow (Proteograph). By limiting the available binding surface of NPs and increasing the binding competition, we identify 1.2 – 1.7x more proteins with only 1% false discovery rate on the surface of each NP, and up to 3x compared to a standard neat plasma proteomics workflow. Moreover, increased competition means proteins are more consistently identified and quantified across replicates, yielding precise quantification and improved coverage of the plasma proteome when using multiple physicochemically distinct NPs. In summary, by optimizing NPs and assay conditions, we capture a larger and more diverse set of proteins, enabling deep proteomic studies at scale.

## 1. Introduction

Blood plasma is an ideal biospecimen to assess human health and disease states because it passes through almost all tissues and is accessible longitudinally and with minimal invasiveness. However, the wide dynamic range of the plasma proteome, over 10 orders of magnitude ^[1,2]^, perhaps millions of proteoforms ^[3,4]^, creates challenges for standard proteomic approaches and prevents wide-spread adoption of untargeted deep proteomics at scale ^[5]^.

To overcome this limitation, we have developed a scalable technology that employs protein-nano interactions ^[6]^. Specifically, introducing a nanoparticle (NP) into a biofluid such as blood plasma leads to the formation of a selective and reproducible protein corona at the nano-bio interface driven by a combination of protein-NP affinity, protein abundance, and protein-protein interactions ^[7,8]^. Remarkably, these nano-bio interactions can be exploited to interrogate the entire plasma proteome at scale and depth without the inherent bias of targeted analyte-specific probes (e.g., antibodies or aptamers). When introduced into a biological matrix, proteins assemble on the NP surface to form a protein corona via physical adsorption and/or electrostatic interactions. Pre-equilibrium, the protein corona composition is based mainly on the relative proximity (i.e., concentration) of proteins that diffuse to interacting moieties on the particle surface. As such, proteins with high abundance dominate the initial corona composition. At equilibrium, governed by thermodynamics, high-abundance low-affinity proteins on the nanoparticle surface are displaced by low-abundance high-affinity proteins (Vroman effect) ^[9]^, which we show leads to compression of the dynamic range. The quantitative composition of protein coronas thus depends on the physicochemical properties of NPs, the presence and abundance of proteins with compatible surface epitopes, and the competition of proteins for NP binding. We previously demonstrated that this process, incorporated within the Proteograph™ proteomics platform, offers superior plasma profiling performance in terms of depth and breadth, compared to conventional workflows ^[6]^.

The competition between proteins for binding surface (Vroman effect) plays an important role in protein corona composition ^[6,9–11]^., and NPs can be tuned with different functionalizations to enhance and differentiate protein selectivity ^[6,9–11]^. In the present study, by modulating the availability of the specific binding surfaces of five different types of functionalized NPs and tightly controlling the ratio of NP to plasma, we modulate the competition of proteins for binding and hence the resulting composition of the protein corona performance of the NP system for deep plasma proteome interrogation. We demonstrate that tuning the Vroman effect by limiting NP surface area significantly enhances downstream proteomics assay performance, improving the coverage of the plasma proteome by as much as 3-fold for a single NP compared to a conventional plasma proteomics (neat) workflow.

## 2. Results

### 2.1. Nanoparticle engineering and experimental setup

To investigate the relation between protein corona composition, physicochemical properties of NPs, and the plasma NP ratio (1-100 fold) (Figure 1A), we engineered, characterized, and compared a series of distinct NPs, comprising three classes with different surface chemical moieties (Figure 1B-E): (1) silica-coated superparamagnetic iron oxide nanoparticles (SPIONs) with core-shell structure and installed carboxylic acid-functionalized silane (NP-2), primary amine-functionalized silane (NP-3), and unfunctionalized surface (NP-1), representing three different types of surfaces of silica-coated SPIONs; (2) SPIONs (NP-4) coated with poly methacrylamide; and (3) glucose-6-phosphate (GSP)-functionalized SPIONs (NP-5). Importantly, the encapsulated SPIONs common to all five types of NPs enable precise and scalable integration into fully automated, magnet-assisted end-to-end sample preparation, from corona formation to protein and peptide isolation for downstream proteome analysis (Figure 1F).

**Figure 1.**
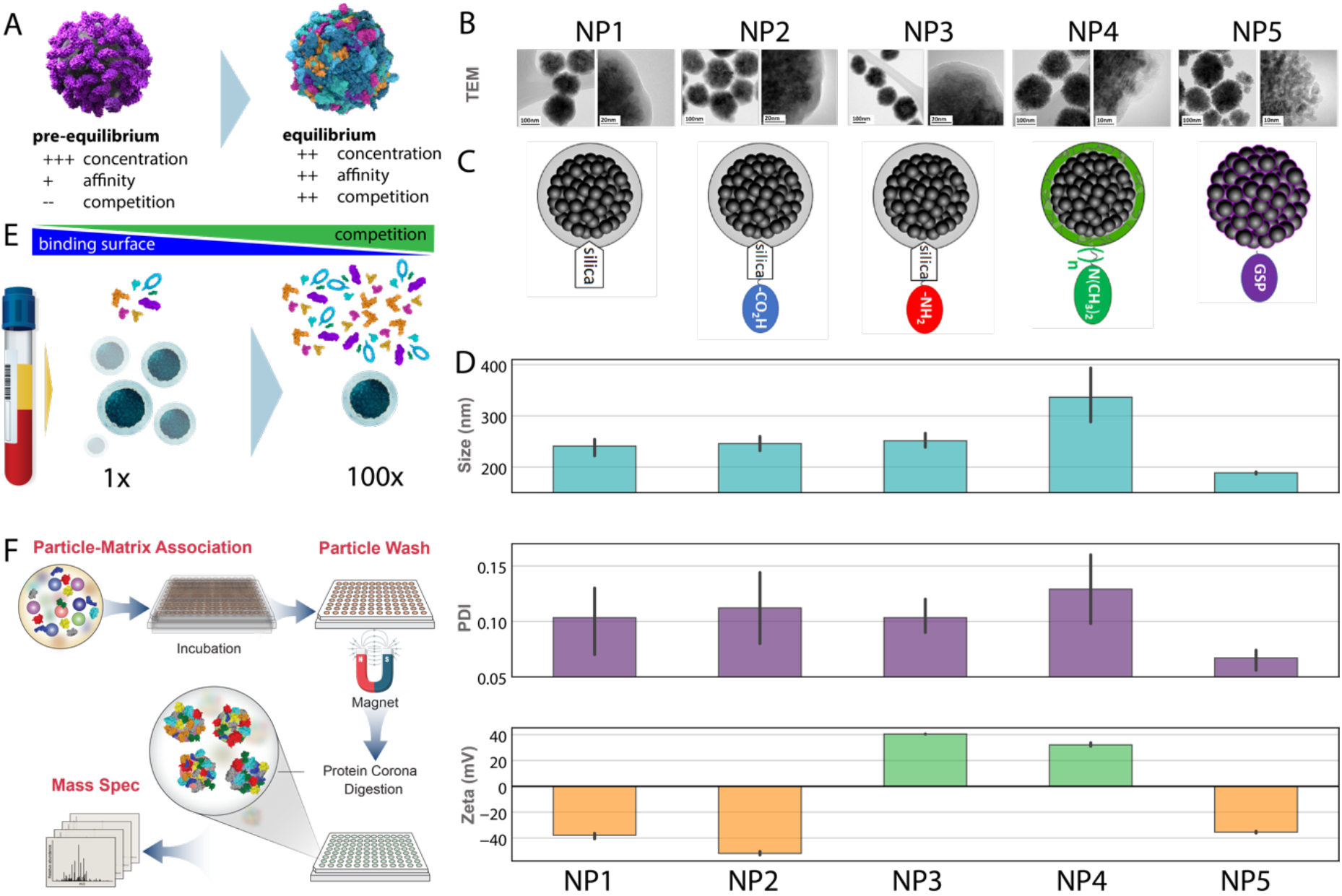
A) Competition drives displacement of high-abundance low-affinity proteins by low-abundance high-affinity proteins. At equilibrium, strong competition for binding increases selective sampling of biomolecules by specific binding affinities, leading to effective ratio compression. B) TEM images C) schematic representation of the NPs. D) Solution-phase properties measured by DLS. Bar plots from top to bottom show the hydrodynamic size (cyan), PDI (purple), and zeta potential (orange and green) of each NP sample in the panel. E) Experimental setup to tune specific nano-bio interactions and competitive replacement (not to scale). F) Proteograph workflow (LC-MS/MS and downstream data processing) schematics.

Transmission electron microscopy (TEM), dynamic light scattering (DLS), and zeta potential measurements were performed to characterize the physiochemical properties of these NPs and to ensure successful synthesis of all 5 target NPs shown in Figure 1B-D. TEM images of NP-1, NP-2, and NP-3 confirm a core-shell structure with an average silica shell thickness of ~14 nm (See Figures 1B). TEM images results of NP-4 and NP-5 show that amorphous polymer layers and small SPIONs (~5-10 nm) are the predominant features present at the surface of these samples, respectively.

Additionally, DLS recorded on aqueous dispersion following synthesis shows that the NPs have hydrodynamic sizes in a range of approximately 200 nm to 350 nm, with polydispersity index (PDI) values varying from 0.05 to 0.15 (Figure 1D). The zeta potential of NPs measured in 5% PBS reveals that NP-3 and NP-4 are positively charged (unlike NP-1), demonstrating that their surfaces were successfully modified with primary and tertiary amine functional groups, respectively. Additionally, a negative zeta potential value was measured for NP-2 and NP-5 NPs, which primarily results from the presence of surface carboxylic acid and phosphate functional groups, respectively. Overall, in-depth characterization including TEM, DLS, and zeta potential measurements confirm the distinct characteristics of the NPs used in this study.

To investigate the extent to which limited surface contributes to NP corona composition and how it can be leveraged to increase utility for deep, untargeted proteome interrogation, we investigated corona composition changes as a function of NP-to-plasma ratios (Figure 1E). Proteins were isolated and processed in a fully automated fashion using the Seer Proteograph ^[6]^. Peptides were analyzed with a trapped ion mobility LC-MS/MS pipeline (timsTOF-Pro ^[12]^) and raw data were processed using DIA-NN ^[13]^ applying 1% FDR cutoff at the protein and peptide levels, yielding 2,482 proteins and 16,270 peptides (2,279 and 14,550 complete identifications in assay replicates, respectively) across the 5 investigated NPs.

### 2.2. Tuning Nano-bio interactions

We investigated the diversity of the protein corona as a function of plasma/NP ratio (P/NP) (Figure 2A). For this experiment, we investigated a single pooled human plasma of deidentified healthy individuals. As effective binding surface area is decreased, the number of proteins identified and quantified (protein IDs) increases. This improvement in individual NP performance compared to neat plasma ranged from 1.8 to 3x (NP-1 and NP-3, respectively), and comparing individual P/NP we found between 1.2x (NP-1) and 1.7x (NP-3) improvements for protein identifications across a P/NP ratio of 1-100x. It should be noted that for NP-1 we excluded one experimental condition that appeared to be an outlier (colored in grey, Figure 2A) from downstream analysis. We next evaluated the extent to which additional protein groups are quantified reproducibly (Figure 2B). Higher P/NP ratios generally yielded more proteins at high precision, with hundreds of proteins being quantified with a coefficient of variation (CV) <10%. Overall, these data demonstrate that NP coronas in general and optimized P/NP proteomics workflows in particular enable deep interrogation of complex biosamples with high precision.

**Figure 2.**
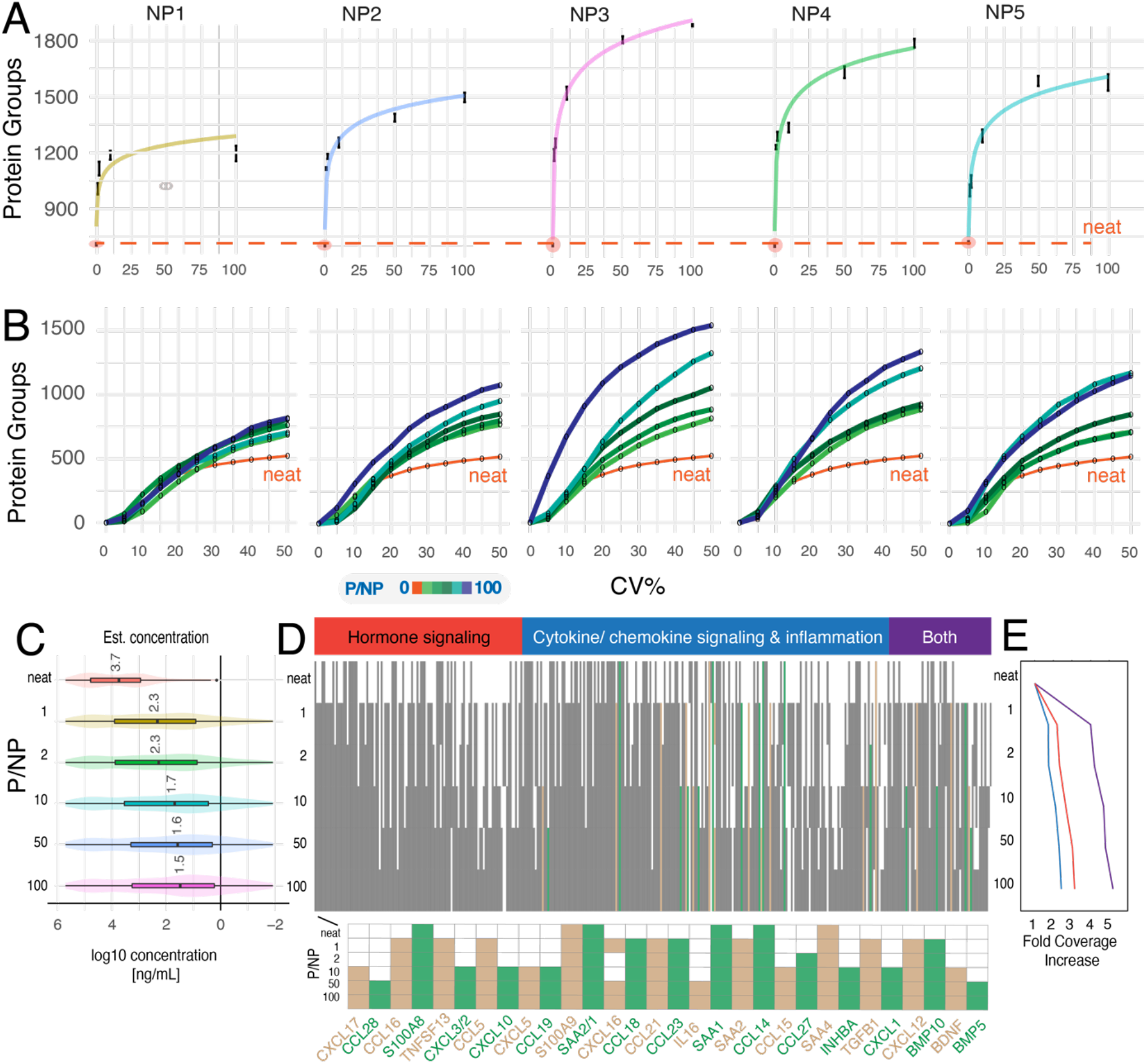
ID, CV, and cytokine/chemokine capture as a function of dilution. A) Number of protein groups at different NP dilution ratios. The error bars show standard deviation across replicates. The P/NP = 0 point shows the baseline for neat plasma. Line shows a regression based on the y ~ a*log(x+0.01) + k. Outliers removed from regression are shown in grey. B) Number of protein groups at different CV thresholds for different NP dilution ratios. Protein groups were filtered for features identified in all replicates (i.e., complete features). Note, outlier samples were removed from analysis. C) Distribution of the concentration of proteins annotated with hormone/cytokine signaling that are present in each NP dilution ratios based the estimated log10 ng/mL values in HPPP reference library ^[14]^. Protein groups were filtered for complete features. Boxplots show 25% (lower hinge), 50%, and 75% quantiles (upper hinge). Whiskers indicate observations equal to or outside hinge ±1.5 * interquartile range (IQR). D) Heatmap showing the presence/absence of genes that have annotations related to hormone signaling, cytokine/chemokine signaling, and inflammation in different NP dilution ratios. Protein groups were filtered for complete features. D) Fold coverage increase compared to neat plasma for each category (hormone signaling in red; cytokine/chemokine signaling in blue; genes involved in both in purple) at each NP dilution ratio. Protein groups were filtered for complete features.

To investigate how increased ID performance translates to better coverage of clinically relevant proteins, we investigated improvement in coverage of functional protein pathways as a function of P/NP. Figures 2C, D, and E show the detection of proteins involved in hormone signaling, cytokine/chemokine signaling, and inflammation based on Uniprot Keywords, GOBP, GOMF, and KEGG terms. The heatmap depicts the increment in coverage of these proteins at panel level (across all NPs) as P/NP increase. The boxplots in figure 2C show that the additional proteins captured at higher P/NP ratios are increasingly lower abundant resulting in deeper interrogation of plasma. In addition, we increase the coverage of each of these groups by 2 – 5 fold when compared to neat plasma. Specifically, chemokines (e.g., CCL14-19, CXCL1, 2, 5), which a standard neat plasma workflow insufficiently captures, were increasingly identified in the biosample under P/NP optimized. Chemokines are an important class of cytokines that exert immune regulatory roles e.g., by guiding migration of immune cells (chemoattraction). Consistent identification of these low-abundance proteins is of high relevance for the study of immune homeostasis and for a variety of acute and chronic diseases. Overall, these results highlight the capacity of tuning P/NP (i.e., Vroman effect) to significantly drive protein identification performance without sacrificing the robustness of corona formation and downstream proteomics readout, facilitating much improved identification of clinically relevant proteins.

### 2.3. Quantitative dissection of nano-bio interactions in complex bio samples

We next investigated individual protein corona changes that drive improvement of protein identification at higher P/NP, focusing on the distinctly functionalized NP-5, NP-4, and NP-2 (Figure 1B-D). Figure 3A shows the delta log10 intensities using the ratio of 1x as baseline. On average, delta log10 intensities increased (boxplots, Figure 3A), demonstrating that the majority of proteins increased in abundance relative to the smallest P/NP. This is consistent with a improved ratio compression, as mean feature intensity improves as well as number of identifications. Importantly, for log-normally distributed protein intensities, reducing the intensity of only a few high-abundance proteins dramatically reduces the signal (i.e., mass of peptides), rendering a large fraction of low-abundance proteins visible to the detector. Consequently, c-means clustering revealed sets of proteins becoming increasingly abundant with higher P/NP (cluster 1, and in part cluster 2, Figure 3B) and sets of proteins with some decreasing intensity trends (cluster 3, Figure 3B).

**Figure 3.**
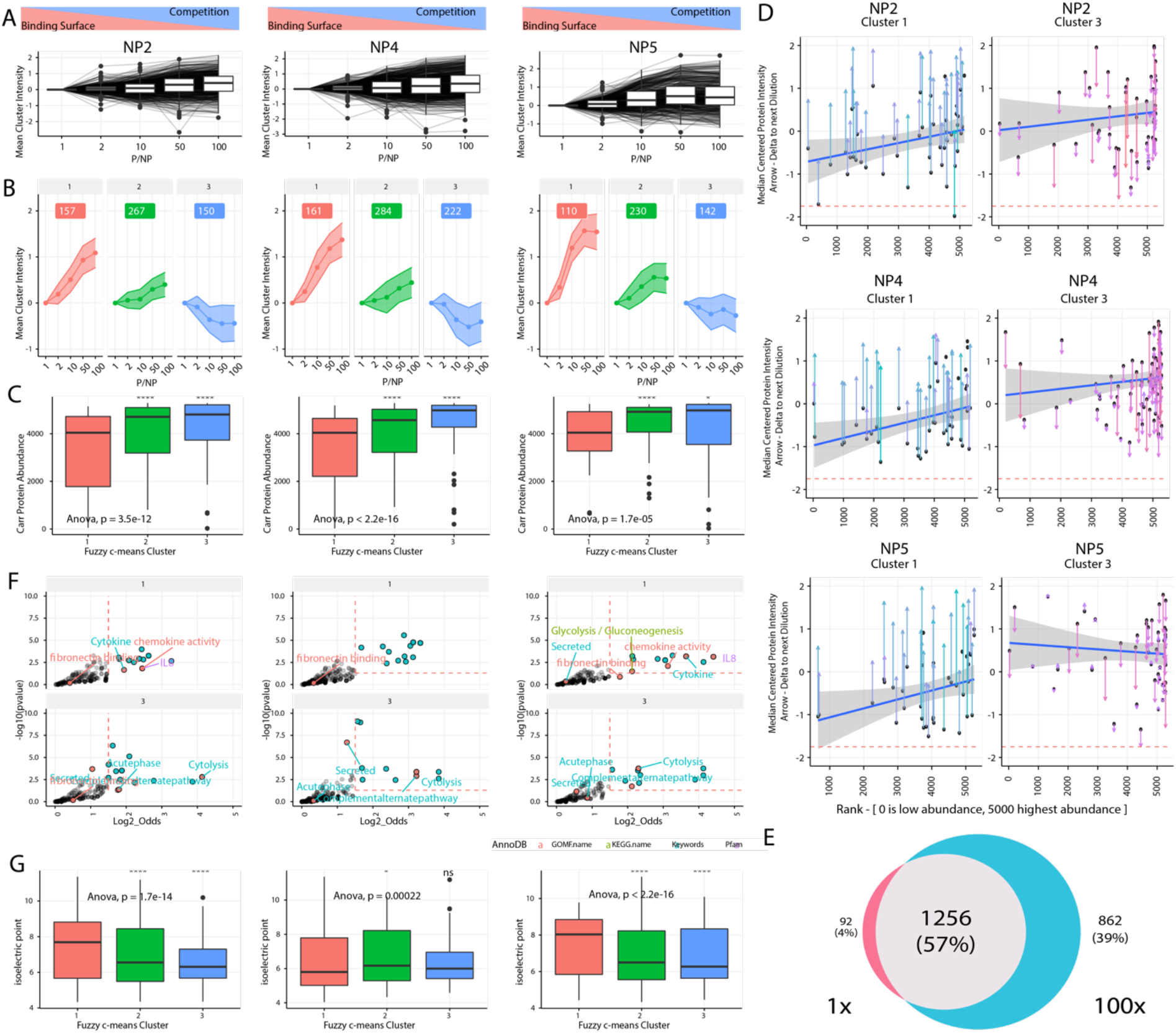
A) Protein group intensity distributions normalized to the lowest (1x) dilution. The boxplots depict an average increase in mean protein group intensities across the dilution series. B) Bootstrapped fuzzy C-means clustering on the protein group intensities using 3 centers, resulting in an increasing (cluster 1), constant (cluster 2), and decreasing (cluster 3) set of proteins across the dilution series. Proteins were filtered (see Method section) prior to that analysis. C) Plasma protein abundance distributions of cluster features. Two-way ANOVA indicates the proteins contained in the three clusters are significantly different. Mann–Whitney U test indicates cluster 1 and cluster 3 contain proteins with significantly different mean plasma abundances. D) Delta intensity comparing P/NP 1x to 100x (y-axis) plotted against rank in a reference database ^[15]^ on the left, proteins with increasing intensities with P/NP ratio (cluster 1) and on the right proteins with decreasing intensities (Cluster 3). E) Number of proteins that are exclusively detected across all 5 NPs at 100x (maximum competition, blue), proteins that are only detected at 1x (minimum competition, red) and the ones commonly detected (grey). F) Enrichment analysis of the increasing and decreasing protein clusters. Upper Row: Over-Representation Analysis on cluster 1 for each NP highlighting Pfam, Uniport keywords, and KEGG Pathway terms with BH corrected p-value < 0.05. Lower Row: over-representation analysis on cluster 3 for each nanoparticle highlighting Pfam, Uniport keywords, and KEGG Pathway terms with BH corrected p-value < 0.05. G) Distribution of isoelectric points for clusters 1-3. C) and G) Two-way ANOVA indicates the proteins contained in all three clusters are significantly different. Welch-test indicates cluster 1 and cluster 3 contain proteins with significant differences in their pI. All boxplots show 25% (lower hinge), 50%, and 75% quantiles (upper hinge). Whiskers indicate observations equal to or outside hinge ±1.5 * interquartile range (IQR). Outliers (beyond 1.5 * IQR) are not plotted.

To further investigate the extent to which signal from high-abundance proteins is attenuated and signal from low-abundance proteins is amplified, we matched protein IDs in clusters 1, 2, and 3 to a reference plasma proteome database ^[15]^. Figure 3C compares approximated true abundance distribution in plasma from the database across the three clusters, confirming that on average lower-abundance proteins are enriched and higher-abundance proteins are depleted at higher P/NP ratios. Figure 3D shows all positive (left) and negative (right) protein corona changes comparing 1x and 100x as a function of their protein rank in the reference plasma proteome database. It shows that a few high-abundance proteins are reduced, improving the signal (i.e., intensity) of a majority of lower-abundance proteins. Across 5 NPs, 862 additional proteins were identified and quantified comparing 1x to 100x (Figure 2E) illustrating the improved performance resulting from the more accessible dynamic range.

We next probed proteins in the two main trend clusters and each NP (up-cluster 1 and down-cluster 3) for common biological roles and functional properties using Fisher’s exact test (Figure 3F), which revealed enriched and depleted protein families within NPs as well as common and distinct trends across NPs. For common trends, the extent of depletion and enrichment is NP specific. For instance, proteins annotated for ‘Complement Pathway’ are depleted (C3) to varying degrees for all NPs. In contrast, the functional groups ‘cytokines’, ‘chemokine activity’, and ‘IL8’ are significantly enriched in C1 for two of the negatively charged NPs (NP-5 and NP-2), are not enriched in the positively charged NP-4, and are slightly depleted in the negatively charged NP-1 and the positive NP-3 (Supplementary Figure S1).

To further explore some of the effects driving corona formation, we investigated the extent to which NPs with negative (NP-5, NP-2) and positive zeta potential (NP-4) increasingly sequester proteins of opposite charge as a function of higher P/NP (Figure 3G). To estimate protein charges at the corona formation pH of 7.4, we predicted the isoelectric point (pI) for each protein. Proteins with a pI > 7.4 are more likely to have a net positive charge at pH 7.4 and *vice versa.* NP-2, which has the most negative zeta potential, showed an enrichment of high pI proteins in C1 and low pI proteins in C3, consistent with an increasing population of positively and negatively charged proteins, respectively. Interestingly, the positively charged NP (NP-4) shows no clear trend of major depletion of positively charged proteins. These results suggest that zeta potential is an important but incomplete descriptor of NPs’ physicochemical characteristics relevant to driving the Vroman effect and protein corona composition in this experiment.

Overall, our data demonstrates that tuning the Vroman effect and competitive binding elicits significant and NP-specific protein corona remodeling, resulting in individual protein and protein functional group and annotation changes. These diversified protein coronas improve ID performance and provide an additional strategy to tailor multi-NP panels for comprehensive proteome interrogation, in addition to changing the functional decoration of the NP surface alone.

### 2.4. Conclusion

Blood plasma is estimated to contain thousands of proteins, and the 22 most-abundant proteins comprise 99% of the overall mass ^[16]^, or in other words, 99% of the raw signal. This skewed dynamic range distribution makes the large majority of relevant biological information undetectable for conventional scalable LC-MS/MS workflows. We have shown previously that nanoparticles (NPs) enable deep and scalable interrogation of the proteome by facilitating a highly reproducible quantitative compression of the proteome’s dynamic range leveraging the Vroman effect. This allows the interrogation of complex biological samples across more than 7 orders of magnitude without the requirement of cumbersome, unscalable workflows entailing high-abundance protein depletion and peptide fractionation ^[6]^. Here we demonstrate that increasing the competition of proteins for binding to NP surfaces substantially improves NP-based proteomics performance in terms of protein identification and precision, resulting in the capture of more-diverse protein coronas that are particularly enriched for low-abundance proteins. This provides an effective strategy in addition to chemical functionalization of NPs to enhance NP-based untargeted proteomics.

In addition to improved protein ID yield by increasingly diversified protein coronas, we observed comparable or higher-precision (CV) quantification and stronger differentiation of protein coronas across functionally distinct NPs. Improved precision could be attributed to better (higher) signal for peptides (resulting in better quantification) but could also be produced by signal (or in this case noise) compression of variation introduced upstream of corona formation. The latter is unlikely, given that most of the steps that could add variance such as proteolysis, solid phase extraction of peptides, or the LC-MS/MS happen downstream of corona formation. Overall these findings suggest that the accumulation of considerably more precisely quantifiable proteins and peptides is the result of improved protein capture, yielding robust signals throughout downstream processing and LC-MS/MS acquisition.

With increased binding competition, low-abundance proteins became more enriched. We saw a strong signature of cytokines and chemokines, which given their low abundance are commonly hard to detect with scalable, unbiased LC-MS/MS workflows. Cytokines and chemokines are involved in a plethora of immunoregulatory processes including immune suppression, immune activation, immune cell chemotaxis; thus consistent identification in blood plasma has high clinical relevance. The unique capacity of NPs to be tuned in terms of their affinity by tailored surface functionalization, combined with enhanced competitive binding, and integrated into a fully automated workflow, therefore provides a powerful and scalable system for biomedical research and discovery.

### 2.5. Experimental Section/Methods

#### 2.5.1. NP Characterization

Dynamic light scattering (DLS) and zeta potential measurements were performed using a Zetasizer Nano ZS from Malvern Instruments. Solutions of NPs (as synthesized) were diluted to a concentration of 5 mg/mL (using 18mΩ water) and sonicated for 10 min prior to testing. Samples were then diluted to approximately 0.02 wt% in DI water and %5 PBS (pH=7.4) for DLS and zeta potential tests, respectively. DLS was performed at 25°C in disposable polystyrene semi-micro cuvettes (from VWR). Zeta potential was also measured using disposable folded capillary cells from Malvern at 25°C following 1 min temperature equilibration time. All reported DLS, PDI, and zeta values are an average of 3 automatic runs performed on each NP sample.

Low- and high-resolution NP imaging was performed using a FEI Tecnai transmission electron microscope (TEM) with an accelerating voltage of 200kV. The TEM grids were prepared by drop-casting 2 μL of NP dispersions (0.1 mg/mL) in water-methanol mixture (25/75 v/v%) on lacey holey grids from Ted Pella, followed by ~24 hours of sample drying in a vacuum desiccator. The shell thicknesses of NPs were measured by image J software through analyzing 50+ individual particles from multiple regions of the TEM grid.

#### 2.5.2. Proteograph Assay

Protein corona preparation and proteomic analysis: NPs were synthesized as described previously ^[6]^. NPs were reconstituted in deionized water to the appropriate concentration for each P/NP ratio ranging from 1x to 100x. The initial concentration (P/NP = 1) for NP1-5 were 60, 170, 60, 40, and 40mg/mL, respectively. Same volume of plasma and NP were mixed together to form the protein corona. The corona formation, wash, protein lysis and alkylation, digestion, and peptide cleanup were done on Proteograph as described previously ^[6]^. After peptide elution, peptide concentration was measured by a quantitative fluorometric peptide assay kit from Thermo Fisher Scientific (Waltham, MA). The peptides were then dried using a Speed Vac. Finally, the dried peptides were reconstituted in the provided reconstitution buffer to a concentration of 0.125 μg/μl.

#### 2.5.3. Data-Independent Acquisition LC-MS/MS

For Data-Independent Acquisition (diaPASEF) ^[17]^ 500 ng of peptides in 4 μL of reconstitution buffer was used (constant mass MS injection). Each sample was subjected to an UltiMate3000 nanoLC system coupled with a Bruker timsTOF Pro mass spectrometer using a trap-and-elute. First, the peptides were loaded on a Acclaim™ PepMap™ 100 C18 (0.3 mm ID x 5 mm) trap column and then separated on a 50cm μPAC™ analytical column (PharmaFluidics, Belgium) at a flow rate of 1 μL/min using a gradient of 5 – 25% solvent B (0.1% FA, 100 % ACN) mixed into solvent A (0.1% FA, 100% water) over 22 min, resulting in a 33 min total run time. The mass spectrometer was operated in diaPASEF mode using ion mobility range of 0.57 – 1.47 V.s/cm^2^ with 100ms accumulation time.

#### 2.5.4. DIA Raw Data Processing

DIA data was processed using the DIA-NN analytical software (version 1.8) in library-free mode ^[18]^. Mass accuracy and MS1 accuracy were set to 10 ppm as it was recommended for diaPASEF datasets. The rest of the parameters were set to default. The FDR cutoff at both precursor and protein level (Lib.Q.Value and Lib.PG.Q.Value) was set to 0.01. Two replicates of the 100 P/NP ratio for NP-3 have been removed from any downstream analysis; one of them was excluded due to issues with the total ion chromatogram and the other one was an outlier in terms of peptide yield compared to the average yield for this condition (7 μg vs. 3 μg).

#### 2.5.5. Data Analysis and Visualization

Data was analyzed in R (v4.0.5) ^[19]^, and visualized with ggplot2 ^[20]^.

##### 2.5.5.1. Clustering

Ratios between the lowest and subsequent dilution median centered protein intensities were computed to generated protein intensities relative to the 1x dilution. These relative intensities were plotted across the dilution series. Ppclust is used to partition data using the Fuzzy C-means clustering. Proteins are partitioned based on bootstrapped I=300 clustering results with n=3 centers. The proteins were assigned to clusters based on maximal average membership probability for each nanoparticle across the dilution series. A minimal cluster probability of 0.5 was used to filter proteins that are not strongly partitioned into any cluster. A clustered median intensity for all proteins in the cluster is shown as a trace with confidence bands showing the mean standard error are computed using R. Median proteins intensity traces within each cluster are plotted for each particle.

##### 2.5.5.2. Enrichment Analysis

Generalized hypergeometric tests for enrichment of Uniprot, Keywords, gene ontology (GOMF) molecular function terms, database of protein families Pfam, and Kyoto encyclopedia of genes, genomes (KEGG) pathways were used for proteins represented in the increasing, constant and decreasing clusters filtered for a Benjamini Hochberg FDR of 5%.

## Data Availability

Data for figure 2, and 3 is available via the PRIDE partner repository ^[21]^ with the dataset PXD030327 (Username; Password: jdrFTjXl). Annotations used for annotation enrichment analysis are available as part of the Perseus framework ^[22]^.

## Supporting Information

**Supplementary Figure S1.**
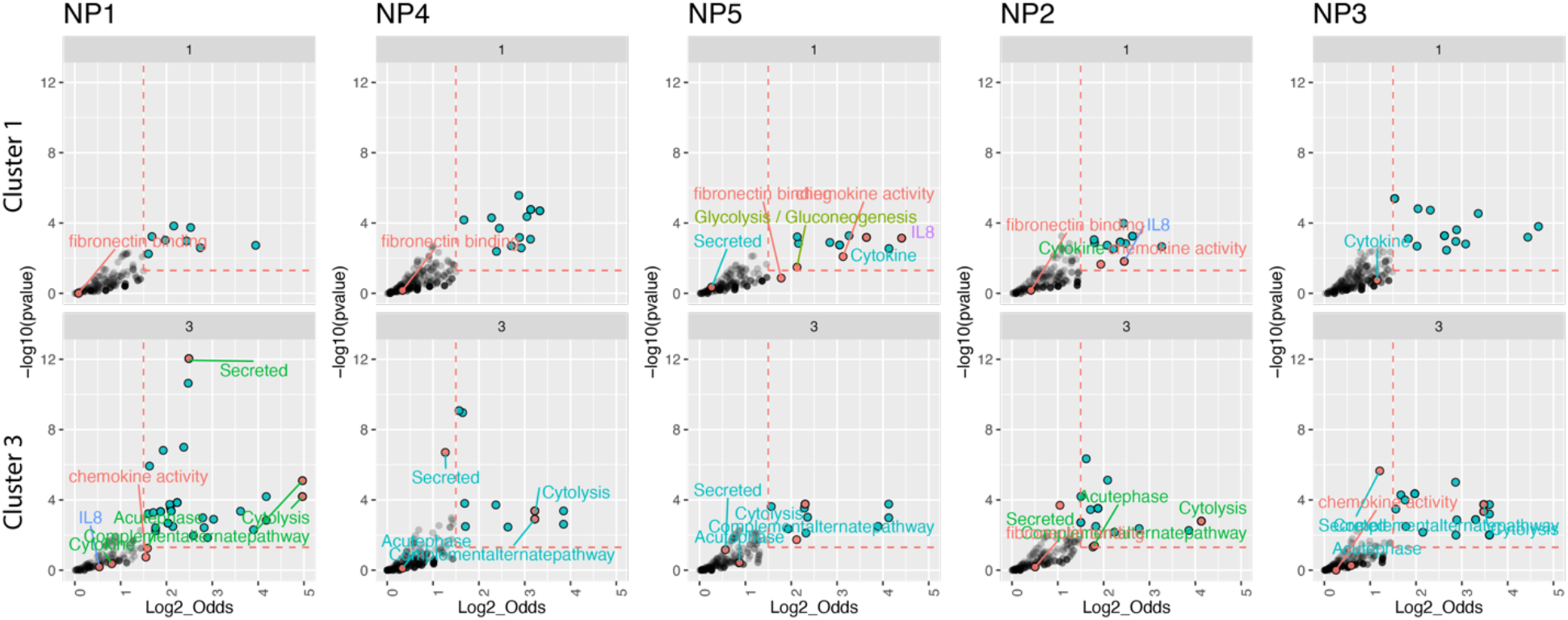
Enrichment analysis of the increasing and decreasing protein clusters. Upper Row: Over-Representation Analysis on cluster 1 for each NP highlighting Pfam, Uniport keywords, and KEGG Pathway terms with BH corrected p-value < 0.05. Lower Row: over-representation analysis on cluster 3 for each nanoparticle highlighting Pfam, Uniport keywords, and KEGG Pathway terms with BH corrected p-value < 0.05.

